# First In Vivo Demonstration of Nose-to-Brain Drug Delivery of Memantine Using NosaPlugs Nasal Inserts

**DOI:** 10.64898/2026.01.26.701702

**Authors:** Emilia Eliasson, Oskar Hallgren, Per-Ola Önnervik, Tomas Deierborg

## Abstract

Treatment of neurological disorders such as Alzheimer’s disease remains a challenge due to ineffective drug delivery to the brain. In recent years, intranasal administration has emerged as a promising non-invasive approach for nose-to-brain delivery. Compared to other routes of administration, nose-to-brain delivery provides a possibility of bypassing both the blood-brain-barrier and the first-pass metabolism in the liver, allowing for a decrease in the delivered dose and thereby a reduced risk of systemic side-effects. While the most common nasal devices, spray pumps, ensure a wide distribution in the nasal cavity and a fast onset of action, a slower release and increased retention time is desired for treatment of many neurological disorders. In this study, we tested the feasibility of a novel nasal insert, NosaPlugs, for prolonged release and delivery of memantine. Using an *in vitro* anatomically realistic nasal model, we demonstrated cumulative release of memantine from the nasal inserts up to eight hours. Additionally, the therapeutic substance was distributed to all parts of the nasal cavity, with higher amounts accumulating in the middle part. *In vivo*, an acute dose of memantine in the gas phase released from the nasal device reached pharmacologically relevant levels in both plasma and the brains of the mice. Future research should investigate the release and delivery of alternative substances interesting for brain diseases, and larger animal models are required to determine the efficacy of nose-to-brain delivery using NosaPlugs nasal inserts. Importantly, our study provides the first proof-of-concept that NosaPlugs can serve as an effective intranasal device for targeted drug delivery to the brain.

## Introduction

Alzheimer’s disease (AD) is the most common form of dementia and is characterized by progressive neurodegeneration and cognitive decline^1^. One of the major challenges in treating AD and other neurological disorders is achieving efficient delivery of therapeutic agents to the central nervous system (CNS). The blood-brain barrier (BBB), a highly selective physical barrier, protects the CNS from potentially harmful substances circulating in the periphery. However, this protective function significantly restricts the physicochemical properties that a drug candidate must possess to successfully penetrate the BBB and reach its target within the brain^2^. Furthermore, systemically delivered drugs must be administered at higher dose levels to achieve therapeutic concentrations within the brain due to low BBB penetration, increasing the risk of systemic adverse events^3^.

Memantine, a non-competitive *N*-methyl-D-aspartate receptor (NMDAR) antagonist, is commonly used to slow the cognitive decline in patients suffering from moderate to severe AD, often in combination with cholinesterase inhibitors^4–6^. By inhibiting NMDA receptors, memantine limits excessive glutamatergic transmission in the brain, thereby reducing symptoms of the disease^7^. The FDA-approved treatment is delivered orally once or twice daily, but due to lack of brain specificity, systemic treatment of memantine is associated with side effects including dizziness, headache, confusion, constipation and diarrhea^8^.

In the last decades, intranasal administration has gained increased attention as a promising non-invasive route for drug delivery to the brain^9^. Apart from being easily accessible, the nasal cavity stands in close anatomical proximity to the olfactory bulb of the CNS. Two distinct pathways, the olfactory and trigeminal, have been identified as routes for direct delivery of therapeutic substances to the brain, bypassing the BBB and first-pass effects^10,11^. Alternatively, substances can be absorbed across the respiratory epithelium and enter the systemic circulation, although this route still requires BBB passage of the substances before reaching the CNS^12^. In these cases, the rich vasculature in the nasal mucosa allows for a rapid absorption of drugs and a fast onset of action, which has been highly advantageous not only for local but also systemic delivery of drugs where a rapid treatment response is desirable, such as migraine^13^ and rescue therapies^14^.

Intranasal drug formulations are predominantly delivered through sprays and aerosols, allowing for an immediate therapeutic effect. However, for certain systemic and neurological treatments a longer retention time is preferred, as well as controlled discontinuation of the drug treatment^9^. The distribution of liquid formulations in the nasal cavity is also highly dependent on the administration technique such as spray orientation and insertion depth, thus increasing the risk for intra- and intersubject variability in dosage^15,16^. To meet the growing interest in nose to brain delivery for drug development, there is a need for innovation of novel intranasal devices that eases patient compliance and allows for a slower and controlled release of therapeutic substances.

Here, we aimed to investigate if a recently developed nasal insert, NosaPlugs, could be used for slow release and efficient delivery of therapeutic substances to the brain. NosaPlugs are silicon-based inserts for nostrils, developed to mask unpleasant external smells without blocking nasal patency. Recently, NosaPlugs have been efficiently used in a study of olfactory memory^17^, and in a clinical trial investigating olfactory training in patients with hyposmia, where the use of scented NosaPlugs resulted in similar treatment response but several-fold higher compliance compared to standard care^18^. In the current study, we first evaluated the rate of memantine release from NosaPlugs and the distribution in the nasal cavity, using an *in vitro* nasal cast model of the human nose. Secondly, we performed a pharmacokinetic study of inhaled memantine in mice following release from NosaPlugs. In brief, we showed that the therapeutic substance could be distributed to all parts of the nasal cavity *in vitro* and released memantine from NosaPlugs was deposited at pharmacologically relevant levels in target tissues *in vivo*. Our study provides the first proof-of-concept that nasal inserts can be used as a device for efficient and prolonged intranasal administration.

## Materials & methods

### NosaPlugs

The nasal plugs tested in this study, NosaPlugs, were developed by NoseOption AB and manufactured by injection molding in their in-house production line. The specific polymer composition and drug-loading method are proprietary to NoseOption AB and cannot be disclosed. However, the loaded dose, release kinetics and performance of the plugs are fully reported below.

### Nasal cast model of the human nasal cavity

A nasal cast model of the human nasal cavity was used for *in vitro* determination of memantine release from NosaPlugs. The cast was 3D printed in polyamide 12 using the selective laser sintering (SLS) technique and consisted of three parts: the front corresponding to the nasal valve and vestibule, the middle corresponding to the turbinates, and the end corresponding to the nasopharynx. All interior surfaces of the nasal cast were coated with a Brij/glycerol solution. For each measurement, ten NosaPlugs containing 20 mg memantine each were placed in the sample chamber, attached to the front part of the nasal cast. A constant airflow of 7.5 l/min was used to release memantine from the plugs. Additionally, three bacterial filters were placed in series and connected to the end part of the nasal cast, to capture any remaining amounts of released memantine not accumulated in the cast. At pre-defined time intervals, the nasal cast and filters were washed with 20% ethanol to extract memantine. Extraction occurred every 30 minutes during the first four hours, followed by extraction every hour up to eight hours, and a final extraction 24 hours after start of the test. Quantification of memantine amounts was performed using liquid chromatography coupled with tandem mass spectrometry (LC-MS/MS).

### Animals

All animal experiments were performed in accordance with Swedish legislation and approved by the local ethical committee (5.8.18-13468/2022). Eight-week-old female Balb/c mice (n=17) were purchased from Janvier Labs and allowed to acclimatize to the new environment for five days after arrival. The mice were housed in groups of 10 mice per cage and maintained at a 12-hour light/dark cycle at constant temperature with free access to food, water and cage enrichment.

### Exposure apparatus

#### Setup of the system

An exposure apparatus (AB FIA, Lund, Sweden) was used to administer gaseous memantine from the nasal inserts (NosaPlugs, NoseOption AB, Stockholm, Sweden) to the mice. The exposure apparatus consisted of two compartments connected with a plastic tube: a delivery chamber for placement of the nasal inserts and an inhalation box containing Batelle holders for each individual animal, ensuring that only the nose was exposed to the drug. Memantine was released in gas-phase from the plugs and distributed into the inhalation box by an airflow, created by pressurized air at the inlet in the delivery chamber and vacuum at two ports in the inhalation box. To measure the concentration of memantine in the inhalation box, 0.22 µm collection filters (Vitalograph) were attached to one of the vacuum ports.

#### Optimization of the delivery system

Prior to the *in vivo* exposure experiments, the delivery system (i.e. the exposure apparatus with nasal plugs) was optimized to maximize the concentration of memantine in the inhalation box and reduce variability.

To optimize the airflow and determine the timepoint of maximum compound release from the nasal plugs, a low airflow (5 L/min of pressurized air and 6 L/min of vacuum) and a high airflow (10 L/min of pressurized air and 11 L/min of vacuum) were tested for 90 minutes each. For this experiment, 10 Nosaplugs containing 30 mg/plug memantine were lined up in the delivery chamber. Two collection filters covered in 1.5 ml of 20% ethanol were attached sequentially to one vacuum port and exchanged every 15 minutes.

To test the variability of the delivery system, the system was tested in triplicate, 30 minutes each run using low airflow, and repeated on a second day. Additionally, separate 30 minutes runs using low airflow were performed to compare the use of 10 or 15 Nosaplugs in the delivery chamber and 30 or 40 mg/plug of memantine, respectively. After each 30-minute run, two collection filters were collected and replaced with new filters covered in 1.5 ml of 20% ethanol. However, on the second day of measurements, three filters were used and collected for analysis.

Memantine was extracted from the collection filters using 10 ml 20% ethanol per filter, and the extracts were analyzed by LC-MS/MS and quantified by the calibration curve.

### Memantine exposure *in vivo*

One day before memantine administration, the mice were acclimatized to the holders in the inhalation box on two occasions for 20 minutes each time.

For the exposure test, 15 plugs containing memantine (30 mg/plug) were placed in the delivery chamber and a low airflow was used (5 L/min of pressurized air and 6 L/min of vacuum). Since the optimization experiments determined that the maximum release of memantine from Nosaplugs occurred between 30 to 60 minutes after airflow onset, the system was allowed to run empty for 30 minutes before animal exposure. All animals were then placed in individual holders and exposed to memantine for 30 minutes. To clear the inhalation box from the active compound at end of exposure (EOE), the vacuum flow alone was left on for one additional minute. Immediately before exposure, the animals had been weighed and allocated into different termination groups. Depending on the termination group, the animals (n=3/group) were anesthetized with isoflurane either at EOE (group 1), 15 min post-EOE (group 2), 30 min post-EOE (group 3), 60 min post-EOE (group 4) or 120 min post-EOE (group 5). As the inhalation box could only hold a maximum of 10 mice at a time, groups 1, 3 and 4 were exposed in one session and groups 2 and 5 were exposed in a second session.

At termination, blood samples were collected by cardiac puncture and mice were perfused with 10 mL PBS. The left-brain hemispheres and left lungs were collected and 100 mg/mL of brain and 50 mg/mL lung tissue per animal were homogenized in distilled water, respectively, using a Bead Mill 24 Homogenizer (Fisher Scientific). Memantine in plasma, lung and brain homogenate samples was quantified by LC-MS/MS.

### LC-MS/MS bioanalysis

#### Preparation of stock solutions and working standards

A 500 µM stock solution of memantine was prepared by dissolving memantine hydrochloride in ethanol. Working standards were prepared by serial dilution of the memantine stock solution in water/ethanol (75:25, v/v) to concentrations corresponding to ten times the desired final concentration in the calibration standards (**Table 1**).

**Table 1.** Concentration of memantine working and calibration standard solutions for bioanalysis.

| Standard | Working solutions diluted in water/ethanol 75:25 (nM) | Calibration standards in spiked plasma, lung or brain homogenate (nM) |
| --- | --- | --- |
| S1 | 40000 | 4000 |
| S2 | 10000 | 1000 |
| S3 | 2000 | 200 |
| S4 | 400 | 40 |
| S5 | 100 | 10 |
| S6 | 40 | 4 |
| S7 | 20 | 2 |
| S8 | 10 | 1 |
| S9 | 5 | 0.5 |
| Blank | 0 | 0 |

#### Preparation of calibration standards and sample solutions

Calibration standards for biological samples were prepared by spiking 54 µl of blank matrix with 6 µl of the working solutions. Mouse CD1 plasma (Sigma-Aldrich) was used as blank matrix for the plasma samples. For lung samples, pooled lung homogenate from three naïve animals were used. For brain samples, brain homogenate from four naïve mice were used as blank matrix. To each 60 µl of calibration standards and biological samples (plasma, lung and brain homogenate), 180 µl acetonitrile was added for precipitation of proteins. Biological samples and standards were thoroughly mixed and centrifuged at 10000 g for 10 minutes at 18 °C. 150 µl of each supernatant was diluted in 450 µl water in LC-vials and vortexed thoroughly prior to analysis.

For filter samples, four calibration standards were prepared by serial diluting the 500 µM memantine stock solution in 20% ethanol to a final concentration of 20, 5, 2 and 0.5 µM. Filter samples extracted in 20% ethanol were transferred to LC-vials for LC-MS/MS analysis. Samples observed to have a higher response than the calibration curve were diluted 1:50 in 20% ethanol.

#### LC-MS/MS conditions

Quantification of memantine in biological and filter samples and in extracts from the nasal cast was performed using a validated LC-MS/MS method. Chromatographic separation was achieved on a Phenomenex Luna® Omega Polar C18 column (1.6 μm, 100Å, 2.1 × 50 mm) maintained at 40°C. The mobile phases consisted of (A) water/methanol/formic acid (95:5:0.1, v/v/v) and (B) water/methanol/formic acid (5:95:0.1, v/v/v). A gradient elusion was applied at a flow rate of 0.3 mL/min, with an injection volume of 2 µL. Mass spectrometric detection was carried out on a Sciex QTRAP® 4500 hybrid triple quadrupole–linear ion trap mass spectrometer equipped with an electrospray ionization (ESI) source operating in positive ion mode. Memantine was optimized for multiple reaction monitoring (MRM) in the mass spectrometer. Data acquisition and processing were performed using Analyst software (version 1.7.3).

Memantine concentration in each sample was quantified based on the calibration curve and the analyte’s absolute peak area. Calibration curves were generated using quadratic regression with matrix-specific weighting: 1/x for plasma samples and 1/x^2^ for lung and brain homogenates and extracted filter samples. The analytical range and lower limit of quantification (LLOQ) of the calibration curves were established based on measured accuracy and precision of the calibration standards. Individual calibration points were accepted and included in the curve if at least 50% of replicate injections had concentrations that did not deviate more than ±20% from nominal values (±30% for the LLOQ). Calibration curves were approved within the concentration range of 1–4000 nM for all matrices.

### Statistical analysis

Differences in body and tissue weights between termination groups were analyzed using one-way ANOVA. Assumptions of normality (Shapiro-Wilk test) and equality of variances (Levene’s test) were assessed prior to analysis, with p>0.05 indicating that assumptions were met. To compare memantine distribution in brain and plasma, a linear mixed-effects model was fitted using restricted maximum likelihood (REML). Fixed effects included time, tissue and their interaction, while mouse was included as a random effect to account for paired tissue measurements from the same animal. When appropriate, pairwise comparisons based on estimated marginal means were conducted. All statistical analyses were performed in R (v4.5.1), using packages dplyr (v1.1.4), car (v3.1-3), emmeans (v2.0.1), lme4 (v1.1-38), and lmerTest (v3.1-3).

## Results

### *In vitro* evaluation of memantine release from the nasal plugs

Memantine release from NosaPlugs (**Figure 1A**) was determined *in vitro* using a nasal cast model and collection filters (**Figure 1B**). Independent measurements occurred for 24 hours, with the highest output rate detected at the second sampling, 60 minutes after start of the experiment (**Figure 1B**). Estimations of the cumulative release of memantine showed that a total of 4987 µg per plug was released over 24 hours (**Figure 1C**).

**Figure 1.**
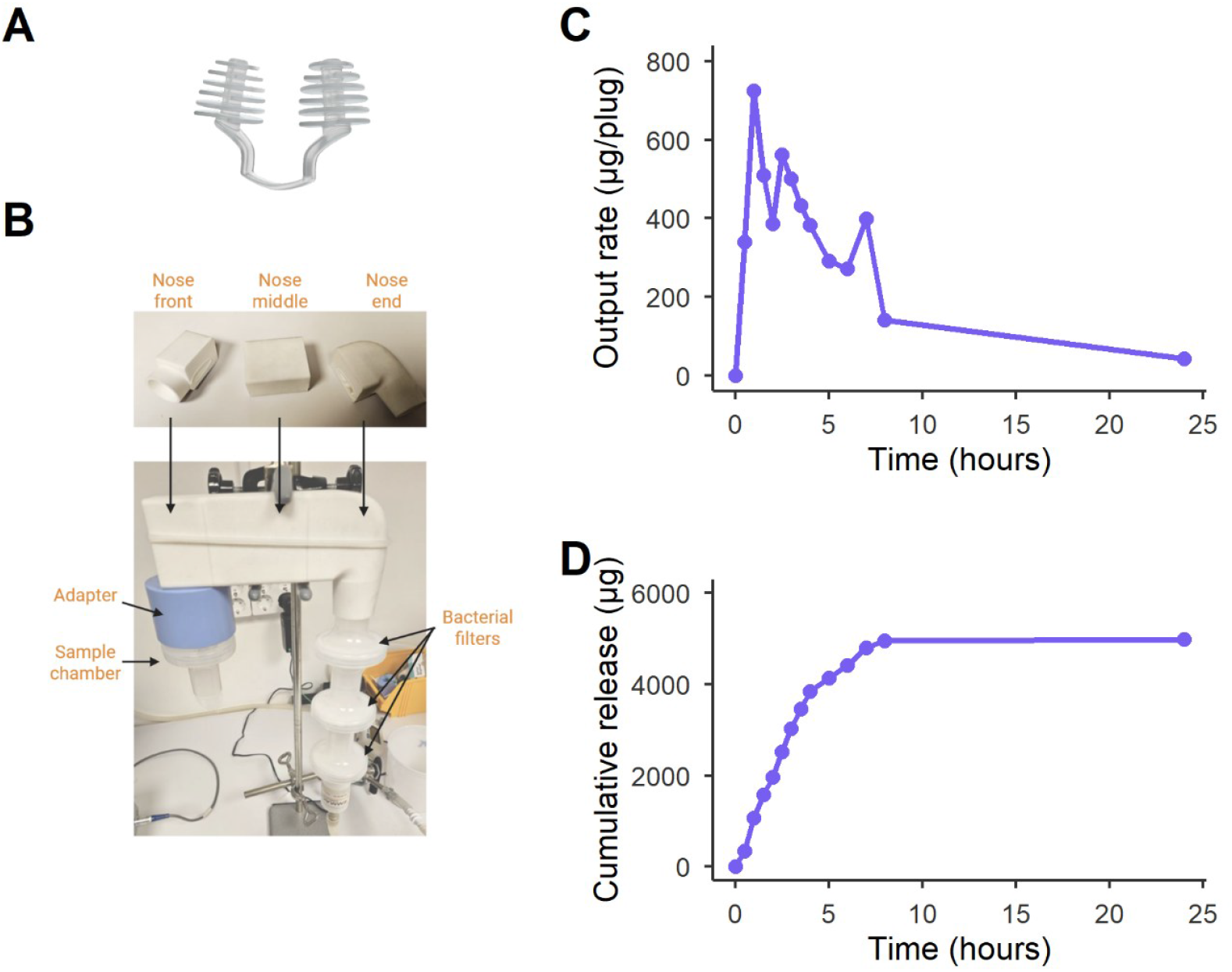
*In vitro* release of memantine from NosaPlugs over 24 hours. Ten NosaPlugs (**A**, 20 mg memantine per plug) were placed upstream of the nasal cast in the sample chamber (**B**). Memantine release at an airflow of 7.5 L/min was measured by LC-MS/MS, which quantified total drug deposition in both the cast and the downstream bacterial filters. (**C**) Output rate per plug. (**D**) Cumulative release.

The distribution pattern of memantine in the nasal cast from each plug showed that 37 ± 13 % (standard deviation, SD) of the total released amount was distributed in the nasal cavity (**Table 2**). The majority of the memantine in the nasal cast, 25 ± 9 % (SD) of total released amount, accumulated in the middle part. A large part of the substance, 54 ± 16 % (SD), was also captured in the first filter but less in the subsequent ones.

**Table 2.** Distribution of memantine (µg/plug) in the nasal cast and collection filters.

| Time (h) | Nose front | Nose middle | Nose end | Filter 1 | Filter 2 | Filter 3 |
| --- | --- | --- | --- | --- | --- | --- |
| 0.5 | 10.33 | 101.56 | 13.45 | 209.10 | 2.32 | 2.58 |
| 1 | 14.76 | 170.69 | 104.97 | 313.22 | 2.37 | 118.63* |
| 1.5 | 101.56 | 174.96 | 12.94 | 216.78 | 2.17 | 1.95 |
| 2 | 15.60 | 134.85 | 11.50 | 221.04 | 1.91 | 2.22 |
| 2.5 | 14.88 | 156.18 | 15.46 | 371.25 | 2.49 | 2.07 |
| 3 | 13.31 | 132.71 | 12.07 | 339.00 | 2.41 | 1.98 |
| 3.5 | 13.93 | 95.16 | 12.89 | 307.24 | 1.89 | 1.32 |
| 4 | 10.92 | 102.41 | 10.72 | 242.00 | 1.89 | 14.71 |
| 5 | 8.60 | 93.03 | 10.92 | 176.66 | 1.64 | 0.97 |
| 6 | 7.73 | 15.02 | 9.61 | 104.12 | 1.17 | 134.85* |
| 7 | 13.31 | 134.42 | 7.89 | 93.88 | 1.07 | 148.07* |
| 8 | 10.57 | 14.29 | 9.01 | 105.40 | 1.19 | 1.02 |
| 24 | 7.94 | 9.46 | 8.77 | 14.53 | 0.98 | 0.98 |

Values marked with asterisks (\*) represent outliers resulting from contamination during sample preparation.

### *In vivo* pharmacokinetic profile of memantine

#### Optimisation of the delivery system

To examine if the nasal plugs could be used to achieve systemic and organ-specific exposure of memantine at pharmacologically relevant levels in mice, an *in vivo* exposure test was performed using an instrumental setup that delivered aerosolised memantine from the plugs to the mice (**Figure 2A-B**).

**Figure 2.**
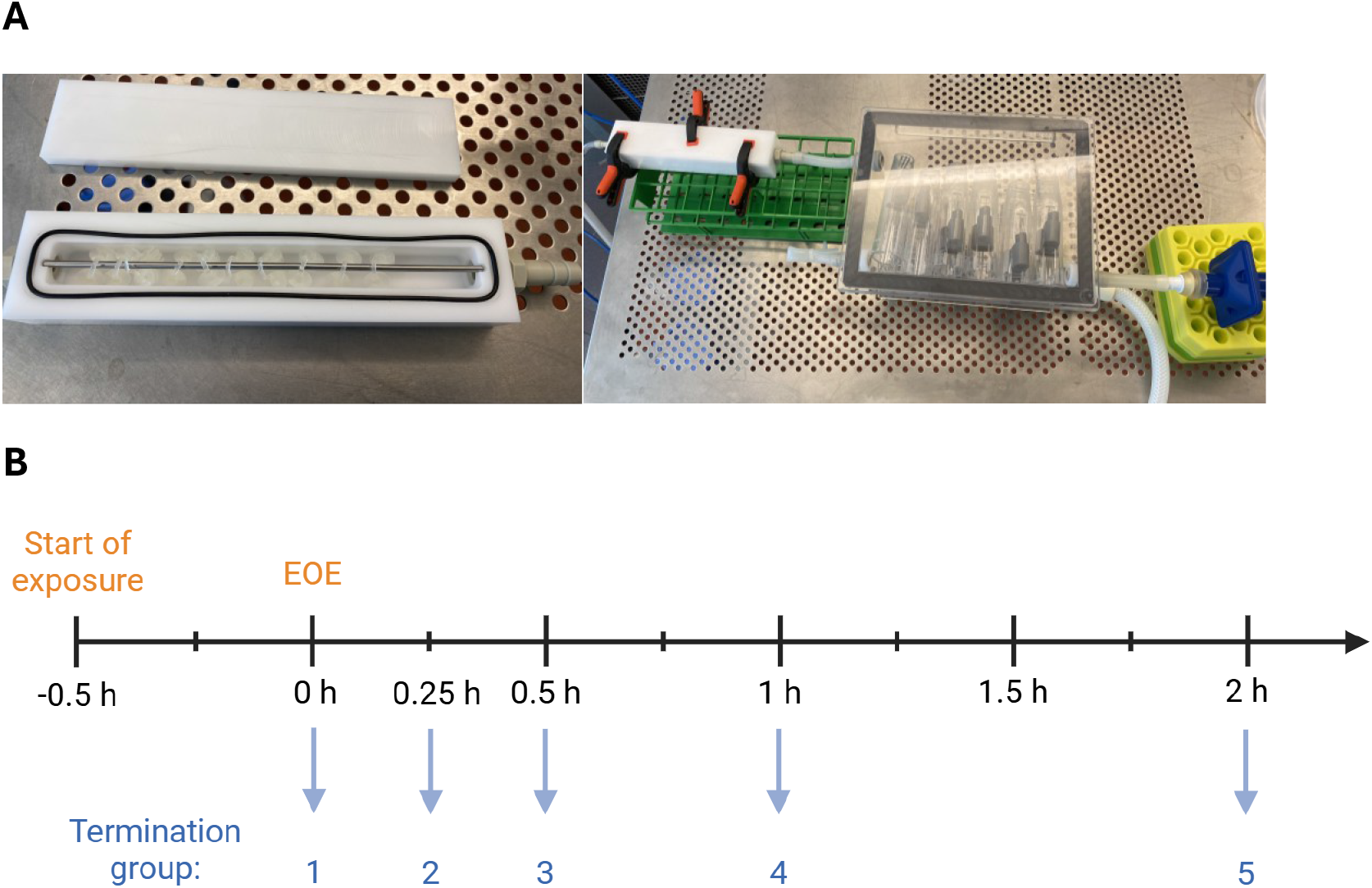
Design of the memantine exposure test *in vivo*. (**A**) Delivery system setup. NosaPlugs containing memantine were lined up in the delivery chamber (left) which was connected to the inhalation box. An airflow generated by pressurized air and vacuum released memantine from the plugs and distributed the substance into the inhalation box where animals were exposed (right). (**B**) Study design. All mice were exposed to aerosolized memantine during 30 minutes until end of exposure (EOE). Depending on the termination group (n=3 mice/group), animals were sacrificed either at EOE, 15 minutes, 30 minutes, 60 minutes or 120 minutes after EOE. Plasma, lung and brain samples were collected from all animals for bioanalysis to determine memantine distribution in target tissues.

To optimize the release of memantine from plugs in the delivery system, a high (10/11 L/min, pressurized air/vacuum) and low (5/6 L/min, pressurized air/vacuum) airflow was tested. The low airflow resulted in higher memantine amounts on collection filters in the inhalation box, with a rapid increase in amount during the first 30 minutes and a first decrease measured at 60 minutes after start of experiment (**Figure 3A**). Increasing the amount of memantine from 30 mg to 40 mg on each plug or the number of NosaPlugs from 10 to 15 in the delivery chamber only resulted in minor differences in released memantine from the plugs (**Figure 3B-C**).

**Figure 3.**
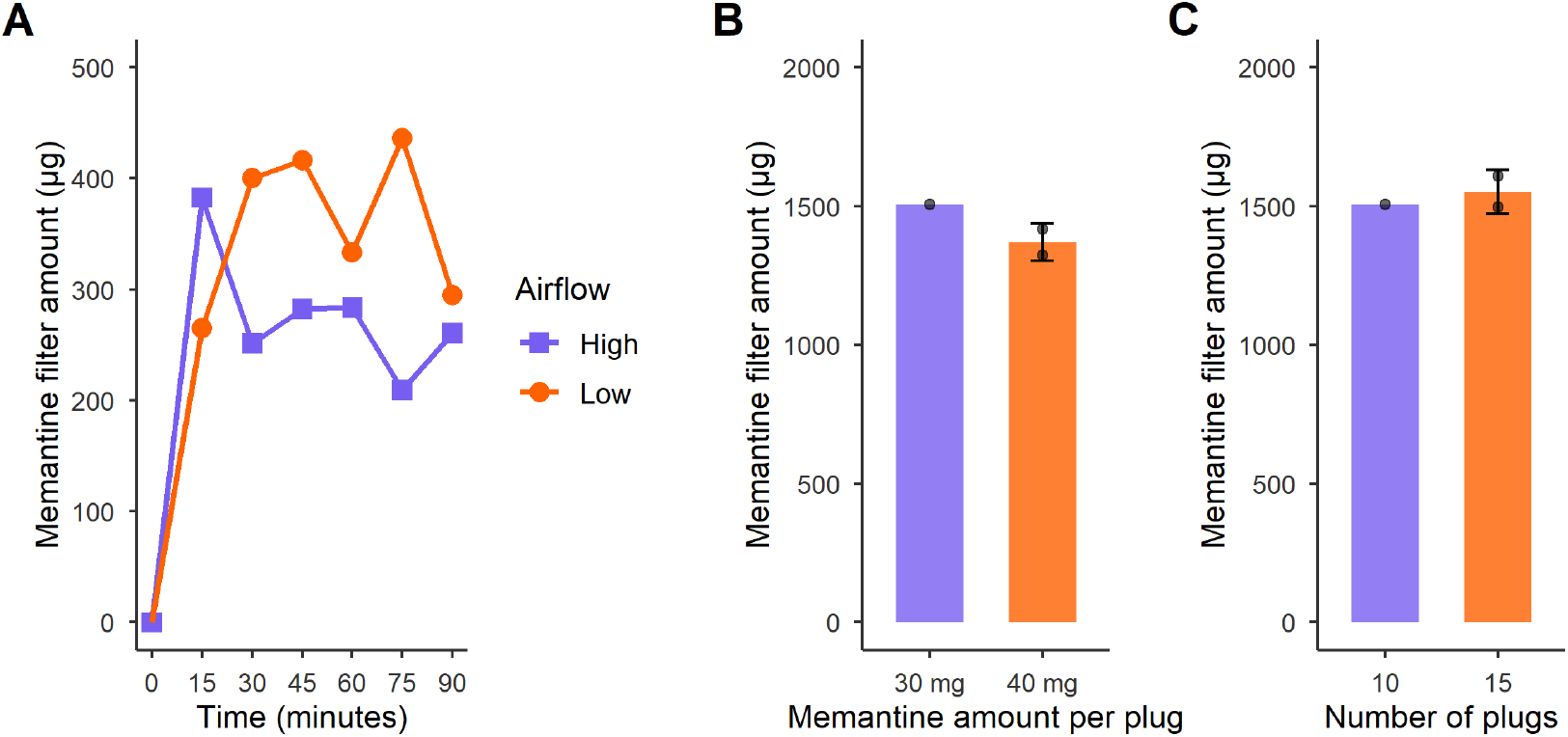
Optimization of the delivery system. NosaPlugs containing memantine were placed in the delivery chamber and substance release from the plugs was estimated based on captured memantine on collection filters in the inhalation box. Filter amounts of memantine were measured using a validated LC-MS/MS method. (**A**) Time-dependent release of memantine from 10 NosaPlugs (30 mg/plug) during a high (10/11 L/min, pressurized air/vacuum) or low airflow (5/6 L/min, pressurized air/vacuum). Comparison of memantine release at low airflow using (**B**) 10 NosaPlugs containing 30 or 40 mg memantine, and (**C**) using 10 or 15 NosaPlugs containing 30 mg memantine in the delivery chamber.

Repeating the experiment for 30 minutes three times using low airflow and ten 30 mg NosaPlugs in the delivery chamber, yielded total memantine filter amounts ranging from 874 to 1018 µg (mean = 923 µg and SD = 82 µg). The non-neglectable amounts of memantine on the second filter indicated that not all substance was captured on the filters, and a third filter was added on the second day of measurements to increase capture (**Figure 4A**). When the experiment was repeated on the second day, total memantine amount of the first two filters reached a similar level as on day one, indicating acceptable intraday and interday variability (**Figure 4B**). However, the amount of substance remained high on the last filter, suggesting that the release of memantine from the plugs may be underestimated.

**Figure 4.**
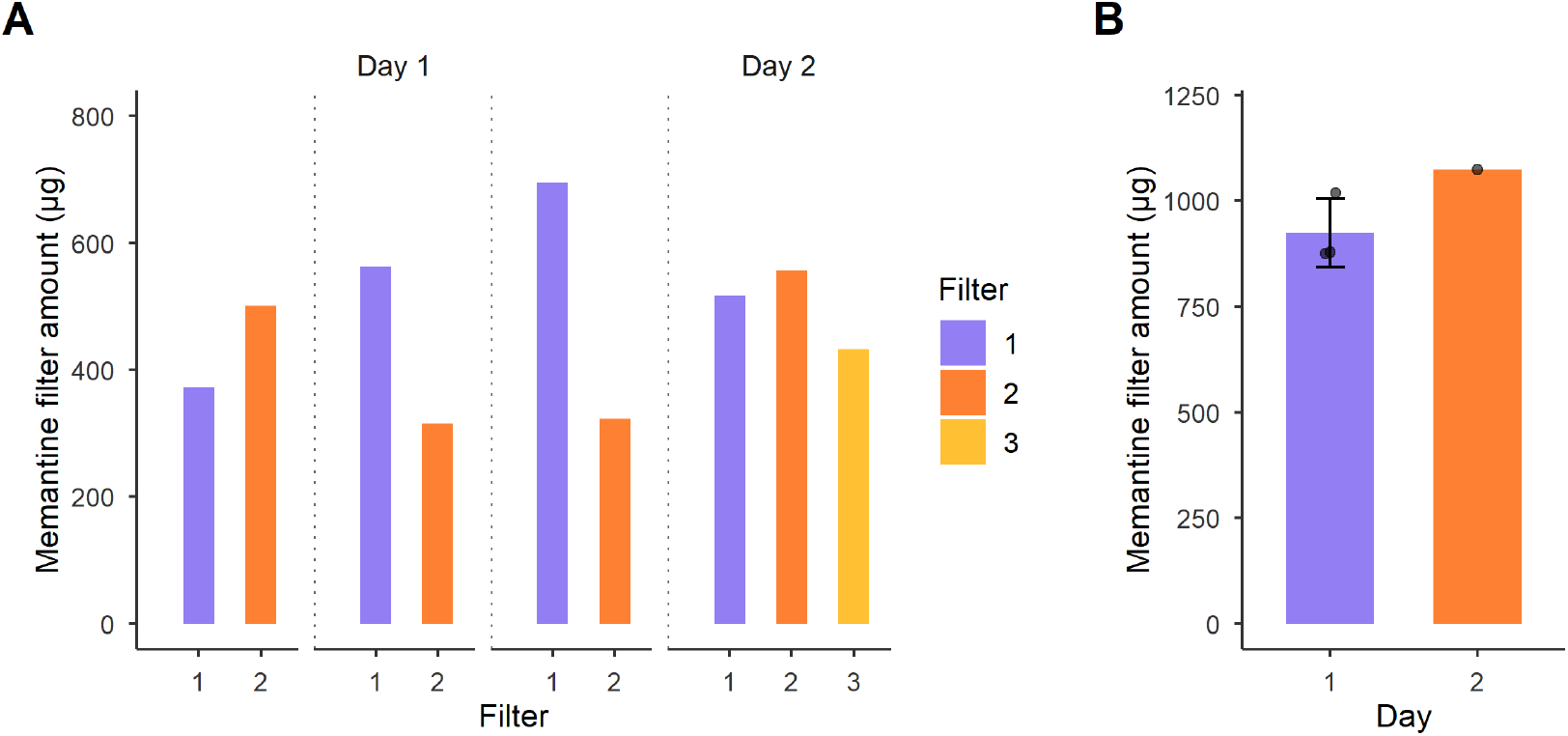
Variability of the delivery system. 10 NosaPlugs containing 30 mg of memantine per plug were placed in the delivery chamber and the system was run at low airflow (5/6 L/min, pressurized air/vacuum) for 30 minutes before the filters were collected. (**A**) Individual filter amounts of memantine. During the first day of measurements, two filters were used to estimate memantine release of NosaPlugs and during the second day, three filters were used. (**B**) Total amount of memantine on the first and second filter during two separate measurement days. On day one, the mean memantine amount was 923 µg (SD = mean +/-82 µg) and on day two, the total memantine amount from filter one and two was 1074 µg.

#### Memantine tolerability

All animals tolerated the memantine exposure test, and no adverse reactions were detected during exposure for any of the termination groups. The body weight and total brain weight were similar between all groups at termination and did not differ from the body weights of the naïve mice (p>0.05 for all groups, **Figure 5A-B**). Additionally, the total lung weight was similar between the groups, despite larger variance between the samples (p>0.05 for all groups, **Figure 5C**). However, due to the small sample size these results should only be regarded as indicatory.

**Figure 5.**
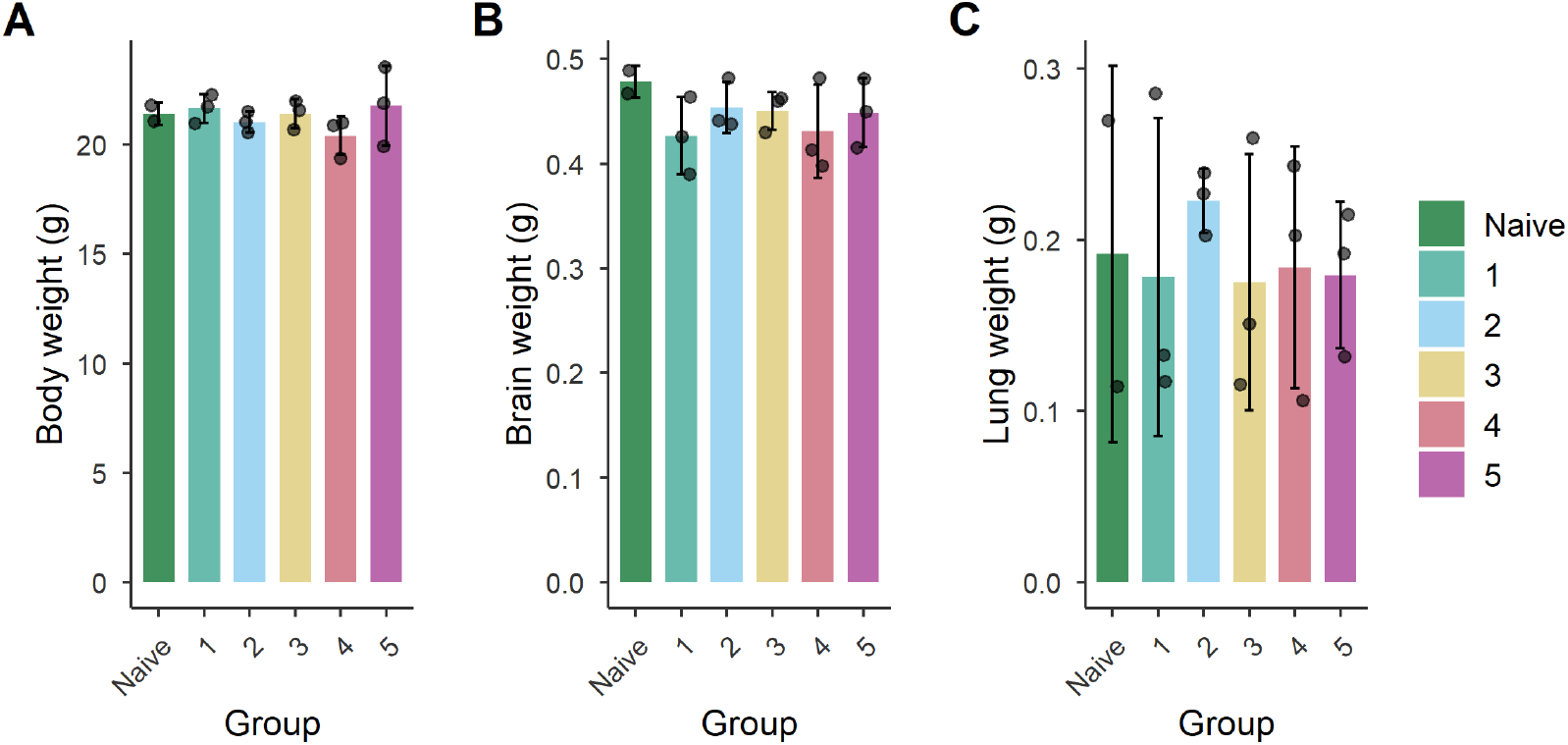
Mean weight of mice and tissue at termination. All mice (n=15) except naïve mice (n=2) were exposed to memantine for 30 minutes through inhalation. Depending on the group allocation, mice were either sacrificed and target tissue was sampled at end of exposure (EOE, group 1), 15 minutes post-EOE (group 2), 30 minutes post-EOE (group 3), 60 minutes post-EOE (group 4) or 120 minutes post-EOE (group 5). (**A**) Mean body weight. (**B**) Total lung weight (left and right lung). (**C**) Total brain weight (left and right hemisphere). Data presented as mean +/-1 SD.

#### Calculation of the delivered dose

To capture and estimate the concentration of memantine in the inhalation box during the inhalation sessions, three bacterial filters were placed in a sequence at the airflow outlet. LC-MS/MS analysis of the filters showed that memantine amounts were similar in all filters irrespective of the order of filters and the inhalation session (**Table 3**).

**Table 3.** Memantine filter amounts following exposure test *in vivo*.

| Inhalation session | Termination groups | Filter amounts (μg) | | | |
| --- | --- | --- | --- | --- | --- |
|  |  | Filter 1 | Filter 2 | Filter 3 | Total |
| 1 | 1, 3, 4 | 455 | 535 | 506 | 1496 |
| 2 | 2, 5 | 697 | 481 | 429 | 1607 |

Based on the filter amounts of memantine, the airflow to the filter (3 L/min), and the duration of exposure, the box concentration of the compound could be calculated (**Equation 1**). Duration of exposure was 30 minutes for both inhalation sessions. Next, the respiratory minute volume (RMV) was calculated (**Equation 2**), using the average animal weight at termination across all termination groups. Lastly, the delivered dose of memantine was estimated based on the calculated box concentration and RMV (**Equation 3**).

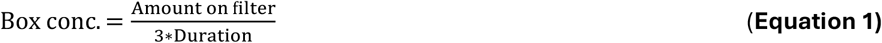

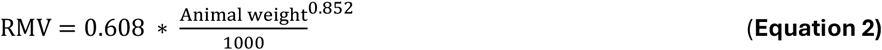

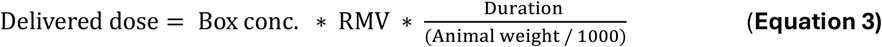

The estimated delivered dose was similar in both inhalation sessions, resulting in an average delivered dose of 0.56 mg/kg (**Table 4**).

**Table 4.** Estimated delivered dose of memantine during each inhalation session.

| Inhalation session | Termination groups | Memantine filter amount (µg) | Animal weight mean (g) | RMV (L/min) | Box conc. (mg/L) | Delivered dose (mg/kg) |
| --- | --- | --- | --- | --- | --- | --- |
| 1 | 1, 3, 4 | 1496 | 21,15 | 0,023 | 0,017 | 0,54 |
| 2 | 2, 5 | 1607 | 21,08 | 0,023 | 0,018 | 0,58 |

#### Tissue concentrations of memantine

Exploratory statistical analysis using a linear mixed-effects model showed that the memantine concentration was significantly higher in brain than in plasma across all timepoints (p<0.001). The highest concentration of memantine in plasma, 1937 ± 197 nM (SD), was detected at EOE. At the last measurement 120 minutes later, the plasma concentration had decreased to 616 ± 28 nM (SD). Meanwhile, memantine concentrations in brain at EOE were 18423 ± 2019 nM (SD) at EOE and a Cmax of 31418 ± 4318 nM (SD) was detected in brain tissue 30 minutes post-EOE. Thus, the brain-plasma ratio increased gradually during the entire 120 minutes, from 9.5 to 33.7 (**Figure 6**). Meanwhile, there was no statistically significant evidence of a time-dependent concentration change (main effect of time) or differences in concentration-time profiles between brain and plasma (time and tissue interaction) during the investigated time interval.

**Figure 6.**
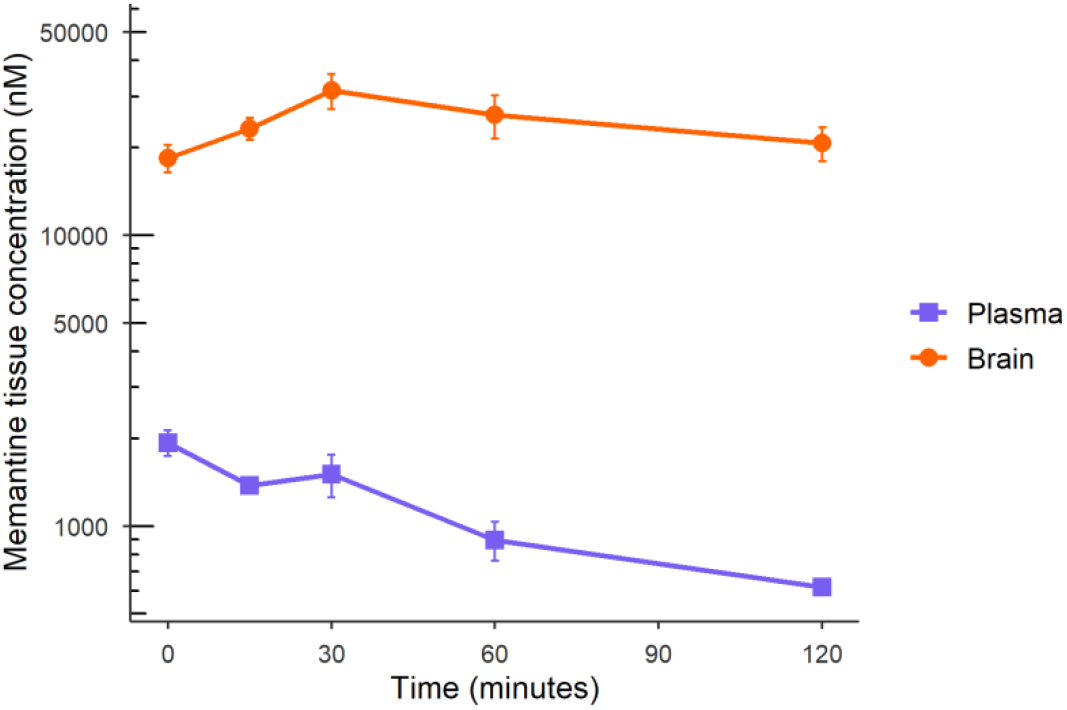
**Pharmacokinetic profile of Memantine** in plasma and brain after 30-minute exposure to memantine released from NosaPlugs. Data presented in lin-log scale as mean +/-1 SD.

The largest within-group variation of memantine concentration was found in lung tissue (**Supplementary Figure 1**). Prior to bioanalysis, brain and lung samples were perfused to remove blood. However, not all lung samples were fully perfused, resulting in blood remaining in the tissue. The highest levels of memantine were detected in the lung homogenate samples that appeared to be visually more contaminated by blood residue (**Table 5**). Thus, these values were excluded from the calculations of the means.

**Table 5.** Memantine concentration in plasma and tissue samples.

| **Animal ID** | **Termination group** | **Administered dose** | **Time (minutes after EOE)** | **Plasma conc (nM)** | **Lung conc (nM)** | **Brain conc (nM)** |
| --- | --- | --- | --- | --- | --- | --- |
| 1 | 1 | 0.54 | 0 | 2070 | 72800* | 20078 |
| 2 | 1 | 0.54 | 0 | 1710 | 58000 | 16174 |
| 3 | 1 | 0.54 | 0 | 2030 | 27000 | 19018 |
| 4 | 3 | 0.54 | 30 | 1240 | 81428* | 27384 |
| 5 | 3 | 0.54 | 30 | 1740 | 67400 | 35973 |
| 6 | 3 | 0.54 | 30 | 1540 | 25800 | 30898 |
| 7 | 4 | 0.54 | 60 | 747 | 17640 | 20803 |
| 8 | 4 | 0.54 | 60 | 935 | 80201* | 28109 |
| 9 | 4 | 0.54 | 60 | 1010 | 36600 | 28779 |
| 10 | 2 | 0.58 | 15 | 1390 | 30600 | 21305 |
| 11 | 2 | 0.58 | 15 | 1450 | 54200 | 25321 |
| 12 | 2 | 0.58 | 15 | 1300 | 28000 | 23090 |
| 13 | 5 | 0.58 | 120 | 592 | 39200* | 23201 |
| 14 | 5 | 0.58 | 120 | 608 | 9905 | 17680 |
| 15 | 5 | 0.58 | 120 | 647 | 19000 | 21305 |
| 17 | naive | 0.00 | 0 | 0 | 0 | 0 |
| 18 | naive | 0.00 | 0 | 0 | 0 | 0 |

Values marked with asterisks (\*) highlights lung samples not fully perfused, thus removed from the calculations of the means. Abbreviations: EOE, end of exposure.

## Discussion

Intranasal administration offers an easily accessible and direct route for drug delivery to the brain. By bypassing the BBB and avoiding hepatic first-pass metabolism, nose-to-brain delivery has the potential to reduce systemic side effects and drug-drug interactions while enhancing local concentrations of the therapeutic agents in the CNS^19^. Thus, nose-to-brain delivery may be especially advantageous for patients suffering from multiple chronic disorders, including the elderly, for whom adverse effects and pharmacological interactions are frequent challenges.

Both preclinical and clinical studies demonstrate that intranasal administration can enhance CNS efficacy relative to systemic delivery for selected therapeutics. Intranasal insulin improves cognitive function without altering peripheral glucose levels^20,21^, and intranasal ketamine produces rapid antidepressant effects and is now clinically approved^22^. In neurodegenerative disease models, intranasal delivery of neuroprotective and anti-inflammatory agents improves behavioral and pathological outcomes, supporting its utility for brain-targeted interventions^23^. For Alzheimer’s and Parkinson’s disease we have found galectin-3 blockage/deletion to be highly efficient to experimentally reduce the pathology^24–27^. Nasal administration of current inhibitors, not passing the BBB, could potentially reach the brain via nasal drug delivery.

In this study, we investigated the feasibility of using a recently developed nasal insert, NosaPlugs, for release and delivery of therapeutic substances targeting the brain. In the first set of experiments, we evaluated the rate of memantine release from NosaPlugs and the distribution in the nasal cavity, using an *in vitro* nasal cast model. We discovered that a third of the released drug was retained within the nasal cavity, the majority of which accumulated in the middle part of the cast. Since this region approximately corresponds to the human turbinates, a region innervated by the trigeminal nerve^28^, it is plausible that the fraction of memantine delivered via the direct nose-to-brain pathway predominantly utilizes the trigeminal nerve rather than the olfactory nerve to reach the brain. However, this hypothesis needs to be further investigated *in vivo*.

Moreover, in our pharmacokinetic study in mice we found that memantine levels in all target tissues were well above 0.5 -1 μM, which is considered pharmacologically relevant^29^. Memantine levels in brain following NosaPlug exposure were approximately 10-35 times higher than in plasma during all measurements, indicating a fast absorption and a good distribution of memantine to the brain following inhalation. These findings are consistent with previous pharmacokinetic studies following subcutaneous, intravenous and per oral routes of administration, which reported brain-to-plasma ratios ranging from approximately 3 to over 20 across all routes^30^. Additionally, the delivered dose of the substance was well-tolerated by the animals, with no clinical observations indicating adverse effects of the nasal drug delivery.

As the NosaPlug device investigated in this study was designed for the human nose, an exposure apparatus was used to deliver aerosolized memantine from the nasal plugs to the mice. Although the deposition dose of memantine in target tissues was measured following inhalation and not specific uptake over the nasal mucosa, the main purpose of this study was to confirm if the NosaPlug device could release relevant amount of the substance to reach sufficient pharmacological levels. Unlike humans, mice are nearly obligate nasal breathers and typically switch to oral breathing only when nasal airflow is obstructed^31^. Consequently, evaporated memantine delivered via inhalation was likely to follow the nasal route, ensuring contact with the nasal mucosa as intended. However, it is important to note that in humans, the nasal plug is also in direct contact with the nasal epithelium, which may increase absorption rate and total exposure to the drug. We also show in our *in vitro* nasal model that NosaPlugs can release memantine to all parts of the cast including the most distant from the airflow inlet, further indicating that aerosolized substances released from NosaPlugs can distribute to the entire nasal cavity.

Although the brain is the primary target of memantine treatment, we found that memantine concentrated at higher levels in lung compared to plasma and brain tissue following 30 minutes of inhalation exposure. Additionally, the high accumulation of memantine in the filters of the *in vitro* nasal model further indicated possible lung deposition of the drug. In mice, the concentration in lung tissue was initially high and remained rather constant during the first 30 minutes after EOE, and after reaching Cmax, the levels steadily decreased and were halved by 120 minutes after EOE. In contrast, we found that levels in brain tissue increased rapidly during the first 30 minutes, followed by a slower decrease during the last 1.5 hours compared to that of lung tissue. To our knowledge, no previous study has reported the pharmacokinetic profile of memantine following inhalation, and only a few have investigated the lung distribution of memantine-like molecules^32,33^. In rats, a high initial accumulation of the compound ZCY-15, synthesized from memantine, was found in lung tissue following IV administration, consistent with our inhalation results^32^. In our study, the highest concentration of memantine was found in lungs that were not perfused completely, which was unexpected as plasma levels were much lower than lung levels. Thus, free memantine in plasma is unlikely to be the sole explanation why higher concentrations of the substance were detected in non-perfused lung tissue. Recent evidence suggests that functional NMDA receptors are present on smooth muscle cells of pulmonary arteries and may be involved in contractility of the vessels^34,35^, which may explain, at least in part, the high accumulation of memantine in lung tissue. Furthermore, the rapid distribution of the substance in lung tissue was likely influenced by the extensive perfusion of the lungs and the high lipophilicity of memantine^33,36^.

Despite promising findings indicating the feasibility of NosaPlugs as a drug delivery platform, some limitations of the current study should be considered. First, during optimization of the *in vivo* delivery system, we found that similar amounts of memantine were detected on the first and second collection filter, suggesting the use of a third filter to increase capture. Although the additional filter increased the total amounts of memantine captured, the substance amounts remained similar in all three filters. These results indicate that the collection filters could not capture all memantine released from the NosaPlugs to the inhalation box, likely causing an underestimation of the delivered dose. To gain a more accurate estimation of the delivered dose, alternative solutions that more efficiently capture evaporated memantine may be needed. Second, as previously mentioned, neither nasal uptake nor mechanisms of blood-brain barrier bypass were investigated in this study. It is noteworthy to mention that memantine is a pharmaceutical with high oral bioavailability in mice and humans^30,37^ and a molecule that readily passes the BBB, in mice presumably through a cationic influx proton antiporter^38^. Thus, we cannot determine if memantine concentrations measured in brain were delivered systemically or through the trigeminal and olfactory nerve pathways. To directly measure nose-to-brain delivery from NosaPlugs, radiolabeling of substances and larger animal models with anatomy more closely resembling humans are needed. Third, only the delivery of one acute dose of memantine was tested here, and the therapeutic effect of the drug following inhalation compared to other routes of administration was not evaluated. However, previous studies investigating the efficacy of intranasal administration of memantine and the cholinesterase inhibitor donepezil in nanocarriers have shown that the nasal route results in higher levels of the drugs in the brain, and increased effects on behavior in mouse models of AD, compared to oral administration^39,40^.

The current study provides a first proof-of-concept that the NosaPlug device enables prolonged release of memantine for at least eight hours and can deliver pharmacologically relevant levels of the compound *in vivo*. These findings underscore the promise of nasal inserts for intranasal drug delivery in neurological disorders, particularly in contexts where a brain-focused release profile is advantageous, and rapid, discontinuous dosing is warranted. To further assess the feasibility of the nasal insert as a medical device for nose-to-brain delivery, additional substances used for treatment of neurological disorders should be investigated *in vivo*. This nasal approach is particularly valuable for drugs with limited blood–brain barrier penetration, which would otherwise be largely inaccessible to the brain.

## Ethics statement

All animal procedures were conducted according to Swedish legislation and by experienced *in vivo* technicians and scientists. Prior to start of experiments, all procedures were evaluated and approved by the local ethical committee (authorization no. 5.8.18-13468/2022). All animals were exposed to maximum one dose of memantine and were monitored throughout the entire study duration. Premature termination would occur for animals reaching the humane endpoints, defined as any signs of pain, complications or behavioral changes, including impaired mobility, altered movement patterns, rough or unkept fur and sitting in a hunched position.

## Author contributions

CRediT: **Emilia Eliasson:** Writing – original draft, Formal analysis, Visualization, Writing – review and editing; **Oskar Hallgren:** Writing – review and editing, Methodology, Resources, Investigation, Formal analysis, Visualization, Conceptualisation; **Per-Ola Önnervik:** Writing – review and editing, Methodology, Resources, Investigation, Formal analysis; **Tomas Deierborg:** Writing – review and editing, Supervision, Project administration, Funding acquisition, Conceptualisation.

## Acknowledgements

We acknowledge the technical expertise of Mårten Svensson, Anna Holmberg, Peter Elfman and Jackie Stuckel at Emmace, who conducted the *in vitro* nasal model experiments.

