## Supplementary Figure 1 for "First In Vivo Demonstration of Nose-to-Brain Drug Delivery of Memantine Using NosaPlugs Nasal Inserts"

### Supplemental material

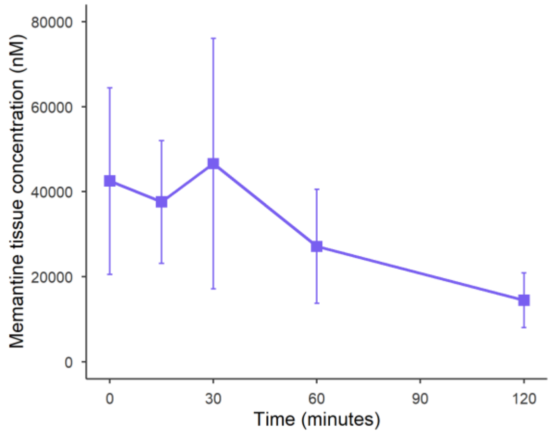


**Supplementary Figure 1. Pharmacokinetic profile of Memantine in murine lung tissue** following 30-minute exposure to memantine released from nasal plugs. Cmax of 46600 ± 29415 nM (SD) was measured 30 minutes post-EOE. Data presented as mean +/- 1 SD.
